# Single-cell molecular subtyping reveals novel intratumor heterogeneity in human Basal-like breast cancer

**DOI:** 10.1101/2024.06.02.597060

**Authors:** Jing Li, Yuan Gao, Shouhui Guo, Qingzhen Hou, Shisong Zhang, Weixing Yu, Ke Liu

## Abstract

Breast cancer is a molecularly heterogeneous disease composed of multiple intrinsic subtypes. Recent studies have highlighted the substantial intratumor heterogeneity of breast cancer, wherein malignant cells of distinct intrinsic subtypes co-exist within the same tumor. However, most existing subtyping methods are designed for bulk transcriptomic data and are therefore limited in their ability to resolve such intratumor heterogeneity at single-cell resolution. To address this gap, we develop UBS93, a computational framework that enables robust molecular subtyping of both bulk tumor samples and individual breast cancer cells. We rigorously validate UBS93 and demonstrate its superior performance relative to existing approaches, particularly in identifying the highly aggressive Claudin-low subtype. Applying UBS93 to single-cell RNA sequencing data from human Basal-like breast cancers, we identify the co-existence of Basal-like and Claudin-low cancer cell populations within the same tumor—a form of intratumor heterogeneity previously observed only in mouse models with genetically engineered RAS pathway alterations. Further analyses suggest that Claudin-low cancer cells originate from Basal-like population, with down-regulation of transcription factor *ELF3* playing a pivotal role in the Basal-like/Claudin-low transition. Together, our findings establish UBS93 as a powerful tool for breast cancer subtyping and uncover a previously unrecognized layer of intratumor heterogeneity in human Basal-like breast cancer.

## Introduction

Breast cancer is a heterogeneous disease characterized by diverse genomic profiles, transcriptomic alterations, and clinical outcomes. This heterogeneity presents significant challenges for achieving clinically meaningful tumor classification^1^. Parker *et al.* utilized microarray technology to develop an intrinsic breast cancer classifier PAM50, which stratifies the disease into five subtypes: LuminalA, LuminalB, HER2-enriched, Basal-like, and Normal-like^2^. This classification system provides superior prognostic and predictive capabilities compared to traditional methods, such as pathological staging, histological grading, standard clinical biomarkers, and so on^2^. In addition to PAM50, Herschkowitz *et al.* identified the highly aggressive Claudin-low subtype which is characterized by low expression levels of cell adhesion molecules (e.g., *CLDN3*, *CLDN4*, *CLDN7*, and *CDH1*) and enriched for high stemness and mesenchymal features^3, 4, 5^. Fougner *et al.* further validated Claudin-low as a distinct subtype through integrative multi-omics analyses across multiple cohorts, although its cellular origin remains unclear^6^.

The use of single-cell omics technology has profoundly enhanced our understanding of intratumor heterogeneity in breast cancer^7-16^. Especially, several recent studies have reported the co-existence of cancer cells of multiple molecular subtypes within the same tumor. For instance, Wu *et al*. identified confident Basal-like cancer cells within a human estrogen receptor-positive (ER⁺) breast tumor (which are usually subtyped as LuminalA or LuminalB)^16^; in addition, Patrick *et al.* demonstrated that modulation of the RAS signaling pathway in mouse models can induce tumors containing both Basal-like and Claudin-low cancer cell populations^5^. These discoveries underscore the limitations of traditional bulk omics-based subtyping strategy and highlight the importance of achieving single-cell resolution for more precise classification.

Despite advances in computational subtyping, significant challenges remain in delineating the intratumoral heterogeneity described above. Most existing tools are optimized for bulk omics data, limiting their applicability in single-cell analyses. To our knowledge, SCSubtype is the only available tool which enables molecular subtyping of breast cancer at single-cell resolution; however, it overlooks the highly aggressive Claudin-low subtype. Addressing these limitations is critical for refining breast cancer subtyping and guiding precise diagnostic and therapeutic interventions.

In this study, we developed a computational framework (UBS93) which enables molecular subtyping of both bulk tumor samples and single cancer cells. Applying UBS93 to single-cell RNA sequencing (scRNA-seq) data, we identified a previously unrecognized form of intratumor heterogeneity in human Basal-like breast cancer, characterized by co-existence of Basal-like and Claudin-low cancer cell populations within the same tumor. Further analysis suggested that Claudin-low cancer cells arise from Basal-like populations, with downregulation of the transcription factor *ELF3* can induce a partial Claudin-low phenotype.

## Results

### Development of the UBS93 classifier for breast cancer subtyping

Solid tumor tissue is a heterogeneous mixture of various cell types^16^. In bulk RNA sequencing, the measured expression of a gene reflects a weighted average of its expression across all constituent cells. Specifically, for a given gene g, if its average expression in cancer cells and non-cancer cells is denoted by C_g_ and M_g,_ respectively, and the tumor purity is α, the observed expression in a bulk sample can be approximated as α * C_g_+(1−α) * M_g._ As a result, gene expression rankings in a bulk tumor sample may deviate substantially from those in pure cancer cells due to the influence of the tumor microenvironment **(Figure 1a, left panel)**; however, for genes highly specific to cancer cells (i.e., M_g_≈0), the rankings in a bulk tumor sample and pure cancer cells are expected to remain highly concordant **(Figure 1a, right panel)**. This observation suggests that the relative expression ranks of cancer cell-specific genes may serve as robust features for molecular subtyping.

**Figure 1:**
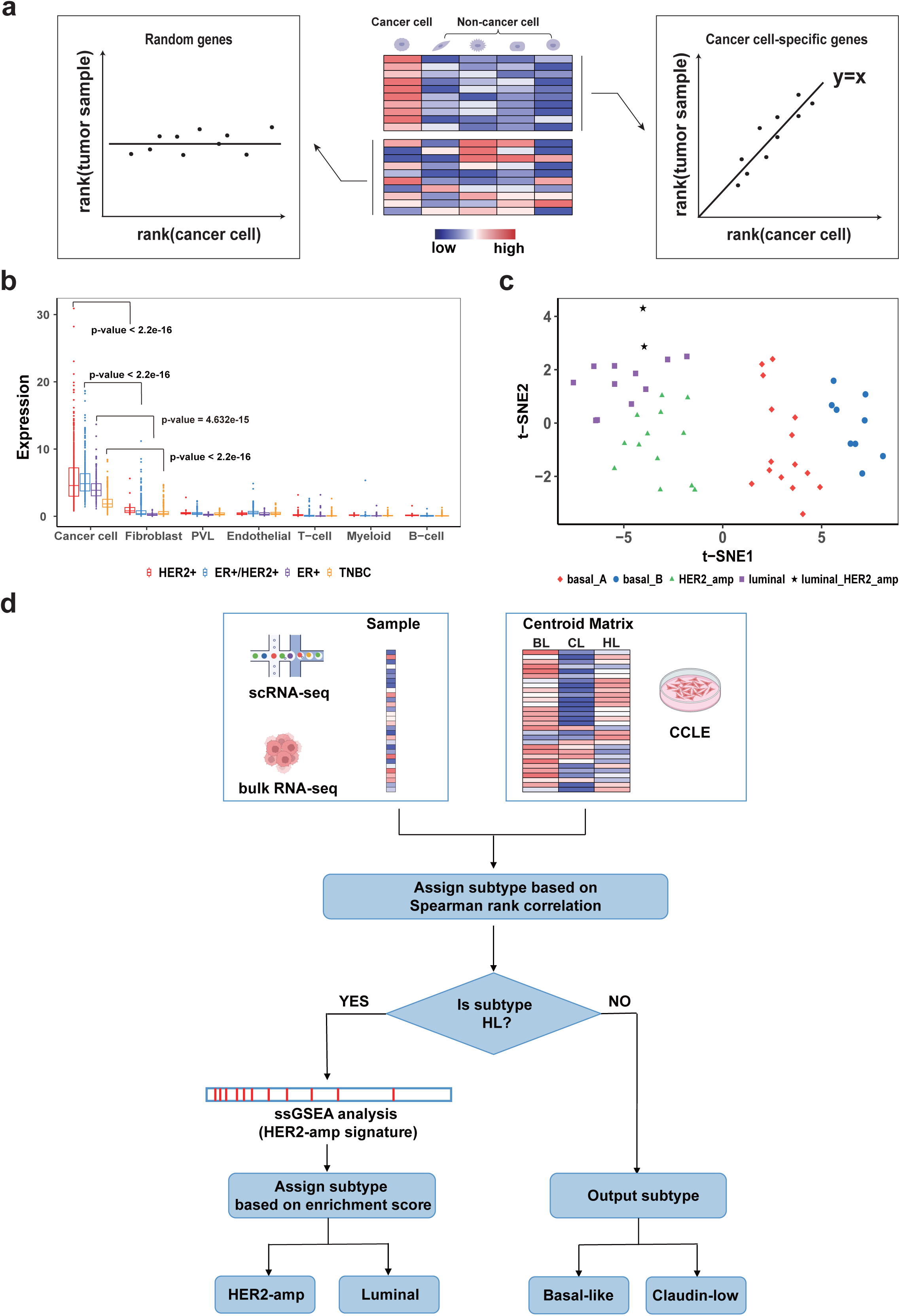
Development of the UBS93 classifier. (a) Schematic representation illustrating the rationale for utilizing cancer cell-specific genes in molecular subtyping. (b) Average expression values of UBS93 genes across various cell types in breast cancer patients. In the boxplots, each dot represents a single cell, the central line indicates the median, and the bounds represent the 25th and 75th percentiles (interquartile range). Whiskers extend to 1.5 times the interquartile range. P-values were calculated using the two-sided Wilcoxon rank-sum test. (c) The t-SNE visualization of CCLE breast cancer cell lines, colored by molecular subtypes. (d) Workflow diagram for the UBS93 classifier. BL: Basal-like, CL: Claudin-low, HL: HER2-amp or Luminal.

To identify such genes, we analyzed the single-cell RNA sequencing (scRNA-seq) dataset from Wu *et al.*’s study and computed "C-M ratio" value for each gene, representing its expression specificity to cancer cells. Among the top 100 genes with the highest C-M ratios, seven lacked ENSEMBL IDs and were excluded. The remaining 93 genes formed a uniform gene panel for breast cancer subtyping, termed UBS93 (**Supplementary Table 1**). As expected, UBS93 genes exhibited significantly higher expression in cancer cells compared to non-cancer cells (**Figure 1b**).

To validate this gene panel, we utilized breast cancer cell line data from the Cancer Cell Line Encyclopedia (CCLE), which classifies cell lines into five subtypes: basal_A, basal_B, luminal, HER2-amplified (HER2-amp), and luminal-HER2-amp^17^. A t-distributed stochastic neighbor embedding (t-SNE) analysis of CCLE bulk RNA-seq data revealed clustering patterns consistent with these subtypes (**Figure 1c**), supporting the validity of the CCLE classification. For consistency with widely used breast cancer nomenclature, we renamed basal_A and basal_B as Basal-like and Claudin-low, respectively, and excluded the luminal-HER2-amp subtype from further analysis. This yielded four refined subtypes: Basal-like, Claudin-low, Luminal, and HER2-amp. We then performed t-SNE using only the UBS93 gene set on the same CCLE dataset (**Supplementary Figure 1**). The resulting subtype-specific clusters mirrored those from the full transcriptome analysis, confirming the utility of UBS93 in distinguishing breast cancer subtypes.

Building on these observations, we developed our UBS93 subtype classifier **(Figure 1d**). We first created a 93 × 3 subtype centroid matrix, where each column represented the average expression profile of UBS93 genes across the CCLE cell lines corresponding to Basal-like, Claudin-low, Luminal or HER2-amp, respectively. For a given bulk RNA-seq or scRNA-seq gene expression profile, we computed its Spearman rank correlation with each subtype centroid and assigned it to the subtype with the highest correlation. If the sample is subtyped as Luminal or HER2-amp, we further calculated an enrichment score based on HER2-amp signature genes to determine the final subtype assignment.

### Evaluation of UBS93 classifier in human and mice

We first evaluated the performance of the UBS93 classifier using an assembled dataset which contains the bulk RNA-seq profile of 47 breast cancer cell lines extracted from the Gene Expression Omnibus (GEO) database^18, 19, 20^**(Supplementary Table 2)**. We found 93.6% of the samples were distributed on the diagonal of the confusion matrix, suggesting the high overall accuracy of our method (**Figure 2a**). Notably, our classifier also demonstrated optimal subtype-specific performance and the F1 scores for the four subtypes were 0.97 (Basal-like), 0.80 (Claudin-low), 1.00 (HER2-amp), and 0.92 (Luminal), respectively. In addition to cell lines, we also evaluated our classifier on the breast cancer samples from The Cancer Genome Atlas (TCGA) and derived ideal overall accuracy (81.9%) and subtype-specific performance^21^ (**Figure 2b**).

**Figure 2:**
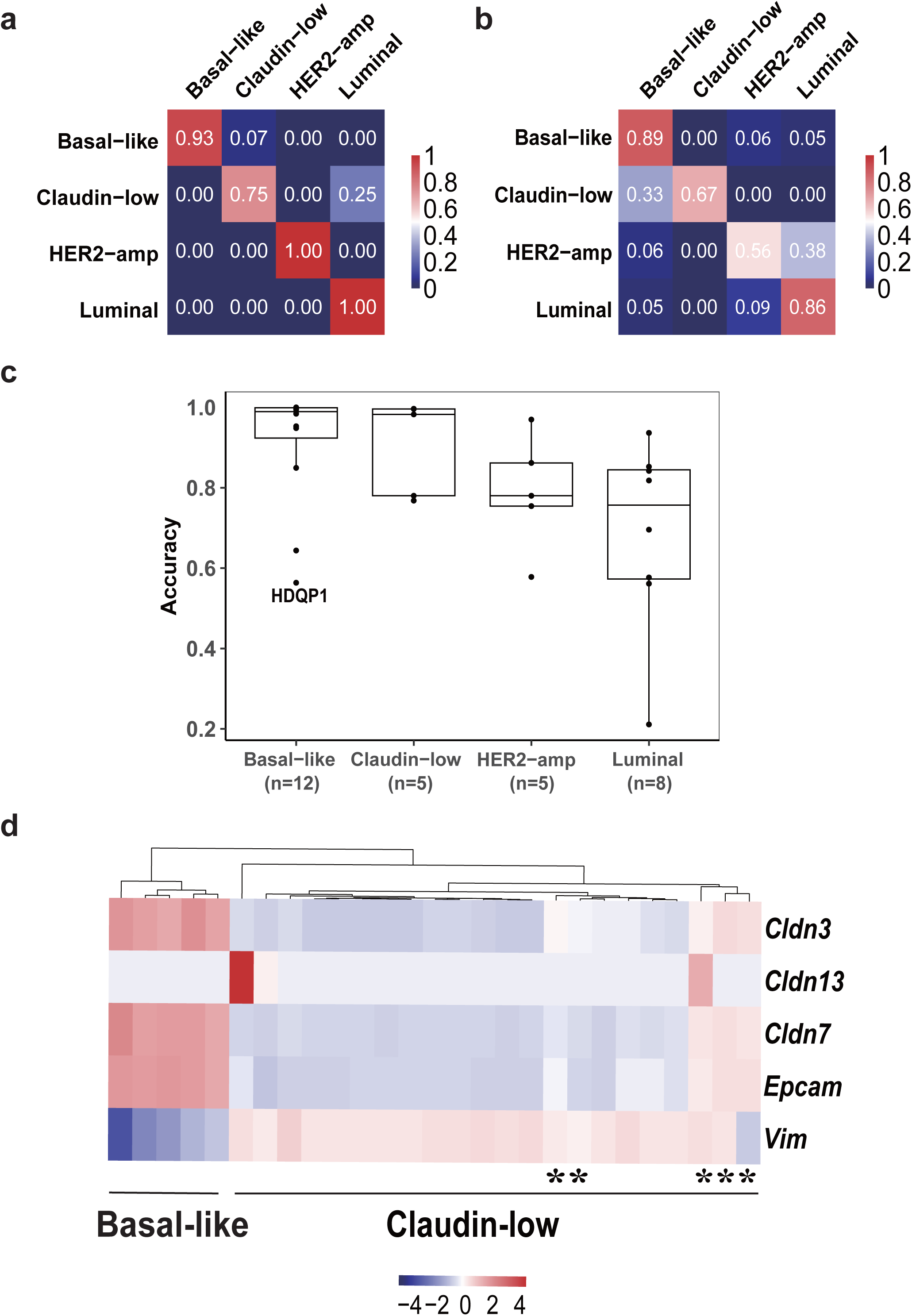
Evaluation of UBS93 classifier in human and mouse. (a) Evaluation of UBS93 using bulk RNA-seq data of breast cancer cell lines. In the confusion matrix, rows represent the actual subtypes, while columns represent the predicted subtypes. In each entry, the value indicates the relative proportion of samples (with the actual subtype corresponding to the row) predicted as the subtype indicated by the column. (b) Evaluation of UBS93 using bulk RNA-seq data from TCGA breast cancer samples. The confusion matrix follows the same format as in (a). (c) Evaluation of UBS93 using scRNA-seq data of breast cancer cell lines. In the boxplots, each dot represents a cell line, the central line indicates the median, and the bounds represent the 25th and 75th percentiles (interquartile range). Whiskers extend to 1.5 times the interquartile range. (d) Scaled expression values of five Claudin-low indicative genes in the assembled murine dataset (27 samples). Asterisks highlight Claudin-low samples whose subtypes were not reported in Rädlers’ study.

Next, we validated our classifier using an assembled scRNA-seq dataset of human breast cancer cell lines (**Supplementary Table 3**)^22, 23^. For each cell line, we computed the accuracy as the proportion of cells that were correctly subtyped. Our classifier achieved an accuracy higher than 80% in 19 of the 30 cell lines (63.3%), with a median accuracy of 85.7%. The subtype-specific median accuracy values were 99.0% for Basal-like, 98.3% for Claudin-low, 78.0% for HER2-amp, and 75.7% for Luminal (**Figure 2c**).

We also validated the classifier on a dataset of 27 bulk RNA-seq samples from murine-derived studies. This dataset included 17 samples from Hollern’s study^24^, which were experimentally confirmed as Claudin-low, and 10 samples from Rädler’s study^5^, with unknown subtype labels. Based on hierarchical clustering of five "Claudin-low indicative" genes (*Cldn3, Cldn13, Cldn7, Vim*, and *Epcam*), we imputed the missing subtype labels (**Figure 2d**). The UBS93 classifier accurately assigned subtypes to all 27 samples, with results aligning with the known subtypes (**Supplementary Table 4**).

### Comparison to existing methods

We systematically compared UBS93 classifier with genefu, a widely used R package for molecular subtyping of breast cancer^25^. Notably, only six genes were shared between UBS93 and PAM50 panels (**Supplementary Figure 2a**); moreover, UBS93 genes exhibited significantly higher C-M-ratio values and cancer cell specificity compared to PAM50 genes (**Supplementary Figure 2b**).

When comparing classifier performance using bulk RNA-seq data from TCGA breast cancer samples, genefu achieved a slightly higher overall accuracy (83.7%). However, UBS93 outperformed genefu in classifying Basal-like, Claudin-low, and HER2-amp subtypes (**Figure 3a**). In murine datasets, genefu misclassified all Basal-like samples as Claudin-low, whereas UBS93 accurately identified them as Basal-like (**Supplementary Table 5**).

**Figure 3:**
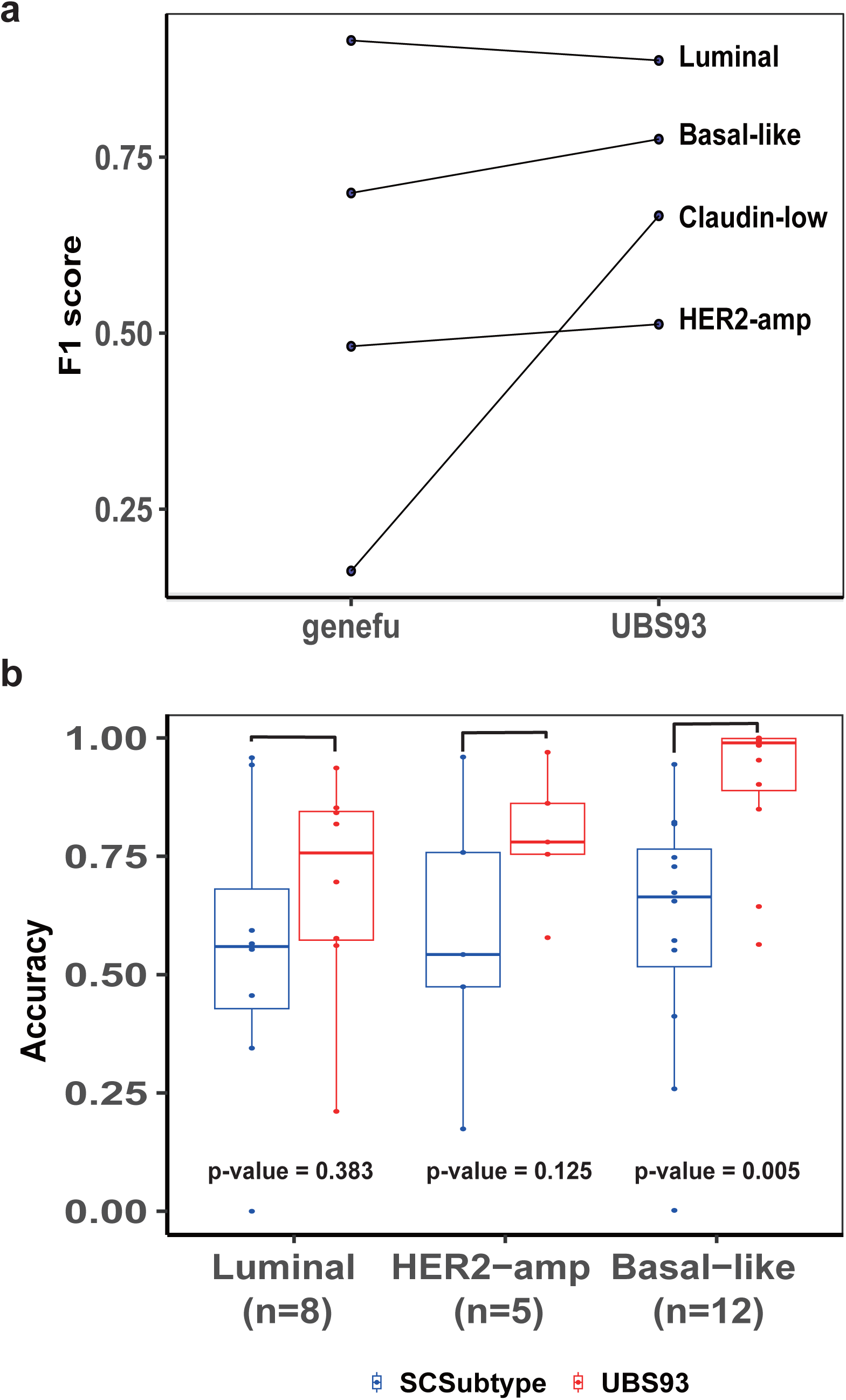
Comparison to existing methods. (a) Comparison of UBS93 with genefu using TCGA breast cancer samples. (b) Comparison of UBS93 with SCSubtype using scRNA-seq data. In the boxplots, each dot represents a cell line, the central line indicates the median, and the bounds represent the 25th and 75th percentiles (interquartile range). Whiskers extend to 1.5 times the interquartile range. P-values were calculated using the two-sided Wilcoxon signed-rank test.

In addition to genefu, we also compared UBS93 with SCSubtype, a method designed for subtyping individual breast cancer cells^16^, using the same scRNA-seq dataset used to evaluate our classifier. Since SCSubtype does not include the Claudin-low subtype, the comparison focused on the other three subtypes. UBS93 demonstrated significantly higher accuracy than SCSubtype for the Basal-like subtype (**Figure 3b**).

### UBS93 uncovers novel intratumor heterogeneity in Basal-like breast cancer

Prior to our study, Wu *et al.* identified Basal-like cancer cells within human ER⁺ breast tumors, highlighting the presence of intratumor heterogeneity in ER⁺ breast cancer^16^. However, the intratumoral heterogeneity of triple-negative breast cancer (TNBC) remains largely underexplored. To investigate this, we re-examined UBS93 subtyping results derived from the scRNA-seq datasets of 12 established Basal-like breast cancer cell lines. In 10 of these lines, over 80% of single cells were classified as Basal-like, consistent with subtyping outcomes derived from bulk RNA-seq data. Strikingly, in the canonical Basal-like cell line HDQP1, only 56.4% of cells were classified as Basal-like, while the remaining cells were subtyped as Claudin-low (**Figure 2c**). These findings led us to hypothesize that, despite being classified as Basal-like based on bulk transcriptomic profile, HDQP1 in fact consists of a mixture of Basal-like and Claudin-low cancer cell populations.

To test this hypothesis, we performed t-SNE visualization of the HDQP1 scRNA-seq data and observed two distinct clusters (**Figure 4a**). One cluster consisted exclusively of Basal-like cells, while 84.2% of the cells in the other cluster were subtyped as Claudin-low. Differential gene expression (DE) analysis suggested that the UBS93-identified Claudin-low cells exhibited canonical features of Claudin-low breast cancer^4, 5, 6, 26, 27^. These included: (1) reduced expression of cell adhesion genes (e.g., *CLDN4*, *CLDN7*, *KRT19,* and *EPCAM*) (**Figure 4b, Supplementary Figure 3, and Supplementary Table 6**); (2) elevated epithelial-to-mesenchymal transition (EMT) level, evidenced by up-regulation of *VIM* and the enrichment of EMT signature genes among up-regulated DE genes (**Figure 4c**); and (3) heightened stemness, as indicated by increased *CD44* (marker of cancer stem cell) expression and higher *CD44*/*CD24* ratio^4, 27, 28^(**Figure 4d**).

**Figure 4:**
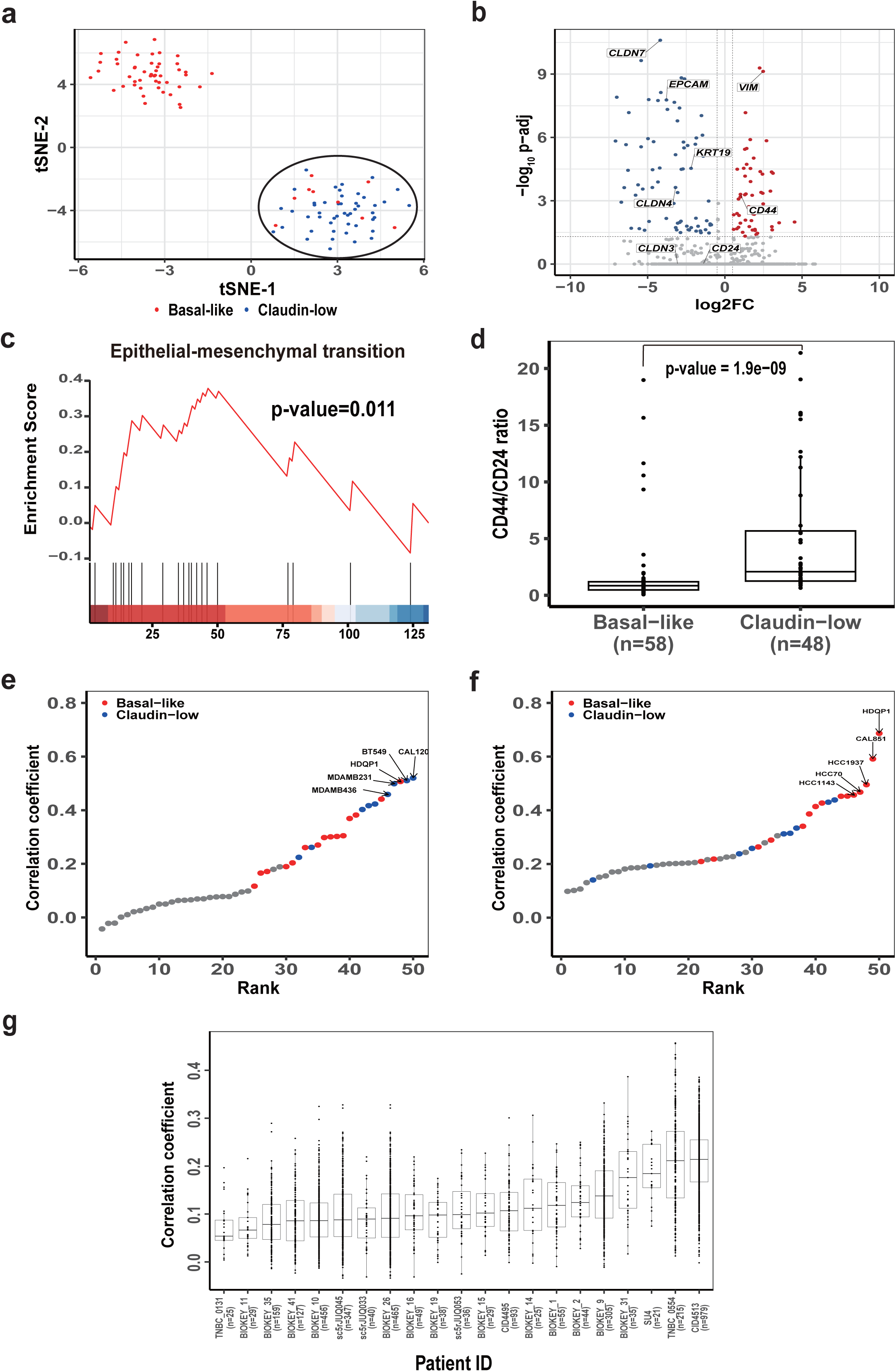
HDQP1 is a cell line with “Basal-like/Claudin-low admixture” phenotype. (a) The t-SNE visualization of HDQP1 scRNA-seq data (GSE17364), with cells colored according to their UBS93 subtypes. (b) Differential gene expression analysis comparing Claudin-low and Basal-like cells of the HDQP1 cell line. Red and blue dots represent genes significantly up-regulated and down-regulated in Claudin-low cells, respectively. (c) Gene set enrichment analysis (GSEA) of epithelial-mesenchymal transition (EMT) signature in the differentially expressed genes. The barcode beneath the heatmap represents the ranked position of EMT genes, from red (high log2FC) to blue (low log2FC). (d) Comparison of stemness between Basal-like and Claudin-low cells. Each dot represents a cell, the central line indicates the median value and the bounds represent the 25th and 75th percentiles (interquartile range). Whiskers extend to 1.5 times the interquartile range. P-values were calculated using the two-sided Wilcoxon rank-sum test. (e-f) Ranking of CCLE breast cancer cell lines based on transcriptomic correlation with Claudin-low (or Basal-like) pseudo-bulk sample. The Basal-like and Claudin-low cell lines are colored red and blue, respectively. (g) Ranking of TNBC patients based on median correlation coefficients with the Claudin-low centroid. Each dot represents a single cell subtyped as Claudin-low by UBS93, the central line indicates the median value and the bounds represent the 25th and 75th percentiles (interquartile range). The whiskers extend to 1.5 times the interquartile range.

To further validate these findings, we generated a pseudo-bulk transcriptomic profile using Claudin-low cells and computed its transcriptomic correlation with breast cancer cell lines from the CCLE project^17^. CAL120—a well-established Claudin-low cell line—exhibited the highest correlation, with four of the top five most correlated lines being Claudin-low **(Figure 4e)**. Conversely, pseudo-bulk analysis of Basal-like cells yielded HDQP1 as the top correlated cell line, with all five top correlates being Basal-like (**Figure 4f**). To double confirm our finding, we replicated our analysis on an independent HDQP1 scRNA-seq data (GSE202771) and derived the same conclusion (**Supplementary Figure 4, 5 and Supplementary Table 7**). Therefore, our computational analysis suggested that the Basal-like cell line HDQP1 was an admixture of Basal-like and Claudin-low cell populations.

To our knowledge, such a “Basal-like/Claudin-low admixture” phenotype has been previously reported only in mouse models following genetic modulation of the RAS signaling pathway^5^, with no prior evidence in human Basal-like breast cancer cell lines and patient samples. To investigate whether this phenotype also exists in TNBC patient tumors, we applied the UBS93 classifier to the scRNA-seq data from 50 TNBC samples. These samples were then ranked based on the median transcriptomic correlation of Claudin-low cells with the Claudin-low reference centroid (**Figure 4g, Supplementary Table 8**). As expected, the metaplastic breast cancer patient CID4513 (from dataset GSE176078) ranked highest, which may not be surprising since previous studies have confirmed that metaplastic breast cancer exhibits characteristics of Claudin-low subtype^29, 30^. Intriguingly, the patient 0554 (from dataset GSE161529), who was classified as Basal-like in the literature, ranked second^15^. Of the cancer cells in this patient, 91.1% were subtyped as Basal-like, while 7.9% were Claudin-low. Similar to HDQP1, Claudin-low cells tended to form a distinct cluster on the t-SNE plot (**Figure 5a**), and meanwhile exhibited canonical features of Claudin-low breast cancer: down-regulation of adhesion genes, enrichment of EMT signatures, and elevated stemness markers (**Figure 5b, 5c, 5d, Supplementary Figure 6, and Supplementary Table 9)**. In addition, pseudo-bulk transcriptomic correlation analysis confirms their resemblance to known Claudin-low breast cancer cell lines (**Figure 5e, 5f**).

**Figure 5:**
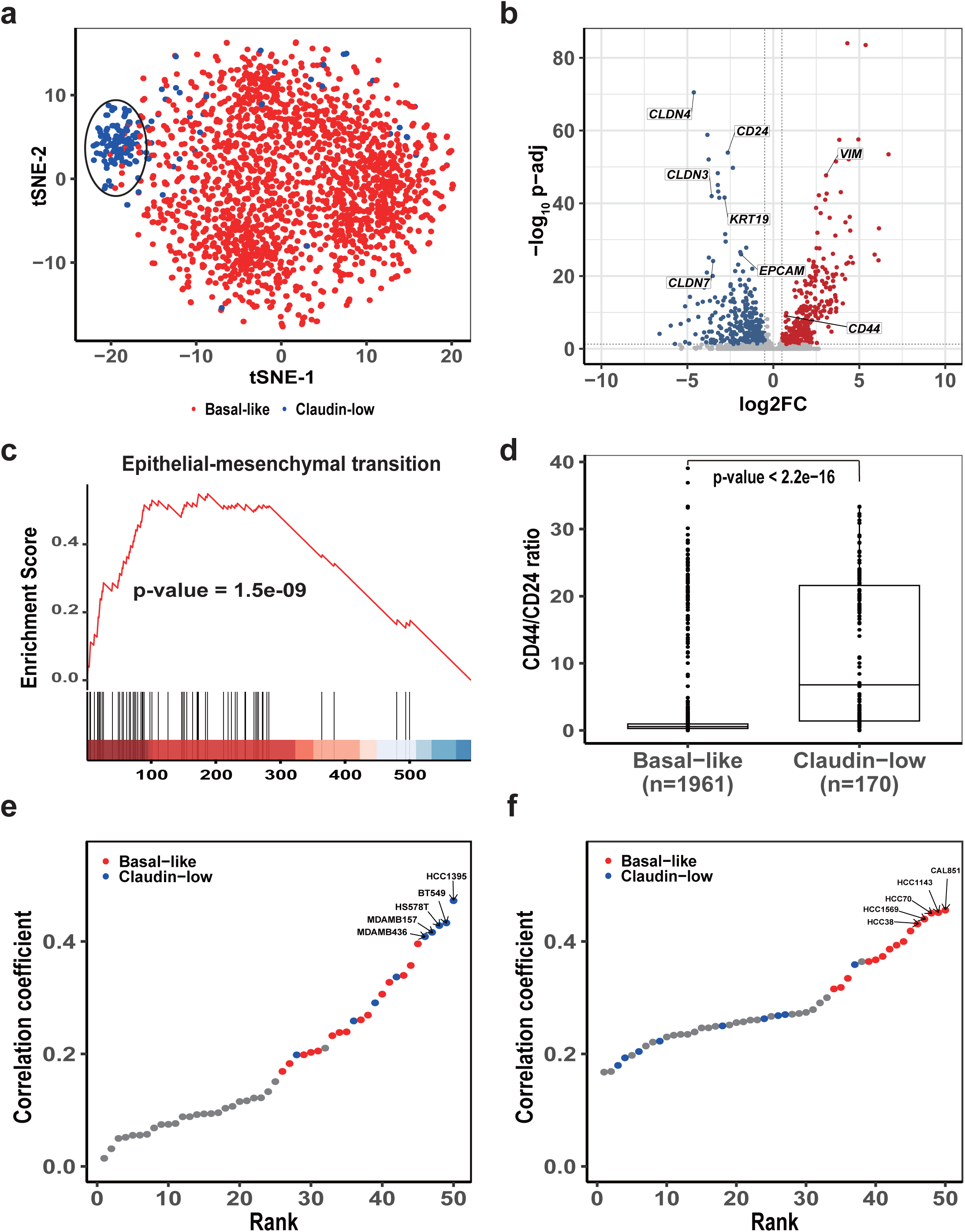
Identification of TNBC patient with “Basal-like/Claudin-low admixture” phenotype. (a) The t-SNE visualization of scRNA-seq data from TNBC patient 0554, with cancer cells colored according to their UBS93 subtypes. (b) Differential gene expression analysis comparing Claudin-low and Basal-like cells in TNBC patient 0554. Red and blue dots represent genes significantly up-regulated and down-regulated in Claudin-low cells, respectively. (c) Gene set enrichment analysis (GSEA) of epithelial-mesenchymal transition (EMT) signature in the differentially expressed genes. The barcode beneath the heatmap represents the ranked position of EMT genes, from red (high log2FC) to blue (low log2FC). (d) Comparison of stemness between Basal-like and Claudin-low cells. Each dot represents a cell, the central line indicates the median value and the bounds represent the 25th and 75th percentiles (interquartile range). The whiskers extend to 1.5 times the interquartile range. P-values were calculated using the two-sided Wilcoxon rank-sum test. (e-f) Ranking of CCLE breast cancer cell lines based on transcriptomic correlation with the Claudin-low (or Basal-like) pseudo-bulk sample. The Basal-like and Claudin-low cell lines are colored red and blue, respectively.

In summary, our integrative analyses reveal a previously unrecognized form of intratumoral heterogeneity in human Basal-like breast cancer, characterized by the co-existence of Basal-like and Claudin-low cancer cells. These findings highlight the capability of UBS93 to detect subtle but biologically meaningful substructures in breast cancer and underscore its potential utility for studying tumor evolution and plasticity.

### *ELF3* knockdown induces a partial Claudin-low phenotype in Basal-like breast cancer

To investigate the underlying cause of the observed heterogeneity, we first examined the cellular origin of Claudin-low cancer cells. Pommier *et al.* conducted large-scale computational analyses of bulk cancer genomics data and proposed three potential origins: direct malignant transformation from normal mammary stem cells, or trans-differentiation from either Luminal or Basal-like cancer cells^32^. However, robust single-cell level evidence from patient samples has been lacking. The identification of Basal-like/Claudin-low admixture in our study provides a unique opportunity to revisit this question. Using copy number variation (CNV)-based hierarchical clustering in patient 0554, we observed that Claudin-low cancer cells were most closely related to Basal-like cancer cells (**Figure 6a**), supporting the hypothesis that Claudin-low cancer cells may arise from a Basal-like lineage.

**Figure 6:**
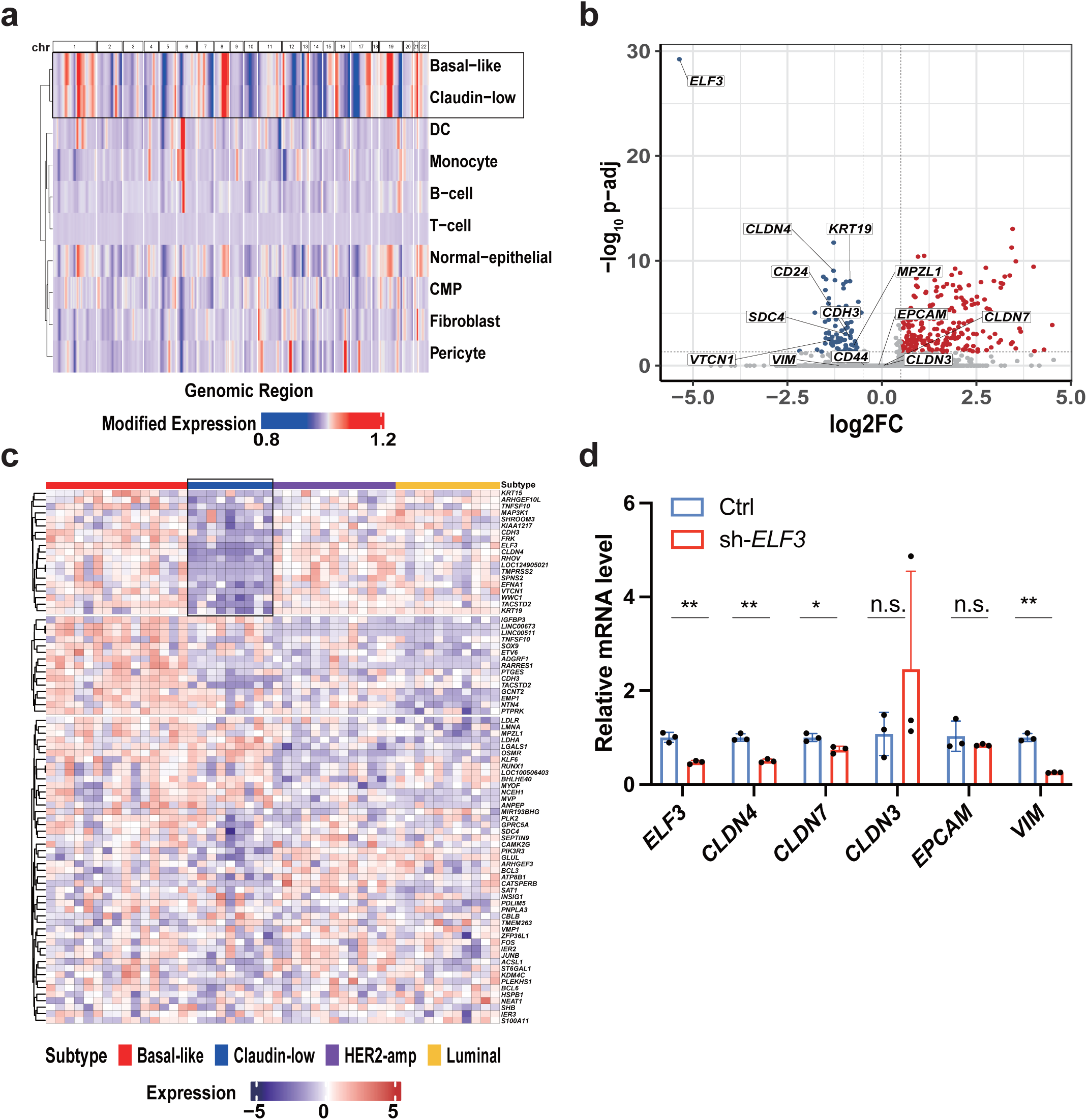
*ELF3* knock-down induces a partial Claudin-low phenotype in Basal-like cancer. (a) Hierarchical clustering of different cell types (from TNBC patient 0554) based on average copy number variation profile. (b) Differential expression analysis comparing sh-*ELF3* and control groups. Red and blue dots represent genes significantly up-regulated and down-regulated in the sh-*ELF3* group, respectively. (c) Scaled expression of down-regulated genes across CCLE breast cancer cell lines of different subtypes. (d) Relative mRNA levels (quantified by RT-PCR) of *ELF3*, *CLDN4*, *CLDN7*, *CLDN3*, *EPCAM*, and *VIM* in control (blue) and sh-*ELF3* (red) groups. Each dot represents a parallel sample. P-values were calculated using an unpaired two-sided t-test. n.s. (p > 0.05), *(p < 0.05), ** (p < 0.01), *** (p < 0.001).

To identify genes which may drive the Basal-like/Claudin-low transition, we intersected DE gene lists from both the HDQP1 cell line and patient 0554, revealing 18 commonly up-regulated and 37 commonly down-regulated genes (**Supplementary Table 10**). Among these, the transcription factor *ELF3* emerged as a gene of interest for two reasons **(Supplementary Figure 7).** First, overexpression of *ELF3* in Claudin-low cancer cell lines (MDA-MB-231 and BT549) inhibits the EMT process^33^; second, *ELF3* promotes *CLDN4* expression in ovarian cancer and non-small cell lung cancer^34,35^. These findings prompted us to test whether depletion of *ELF3* in Basal-like cancer cells might induce a Claudin-low phenotype.

We selected HCC70, a cell line with a transcriptomic profile closely resembling Basal-like breast cancer³⁶, as our experimental model. scRNA-seq followed by UBS93 subtyping confirmed that 98.7% of untreated HCC70 cells were classified as Basal-like. We then performed *ELF3* knockdown in HCC70 and repeated scRNA-seq to determine whether this perturbation could promote Claudin-low features. UBS93 analysis showed that 98.1% of cells remained Basal-like, suggesting that *ELF3* knockdown alone is insufficient to trigger a complete subtype switch.

Despite failing to induce complete Basal-like/Claudin-low transition, we asked whether *ELF3* knockdown could induce partial Claudin-low features. To account for variable knockdown efficiency, we selected the 100 cells with the lowest *ELF3* expression (sh-*ELF3* group) and the 100 cells with the highest *ELF3* expression (control group) for further analysis. Interestingly, the sh-*ELF3* group exhibited a significantly lower correlation with the Basal-like centroid compared to the control group (**Supplementary Figure 8a**) and DE analysis between these two groups identified 210 up-regulated and 79 down-regulated genes, respectively (**Figure 6b, Supplementary Table 11**). As anticipated, *CLDN4* was significantly down-regulated (p-value = 3.57e-14), as was the epithelial cell marker *KRT19* (p-value = 3.59e-13). Interestingly, when examining the expression pattern of down-regulated DE genes across CCLE breast cancer cell lines, several exhibited Claudin-low specific depletion (**Figure 6c**). We curated a set of such genes (which we termed as CLSD genes) and gene set enrichment analysis revealed significant enrichment of CLSD genes among the down-regulated DE genes (**Supplementary Figure 8b**).

In addition to scRNA-seq, we also used qRT-PCR to assess how *ELF3* knockdown influenced the expression of *CLDN3*, *CLDN4*, *CLDN7*, *EPCAM*, and *VIM* (**Figure 6d**). Consistent with scRNA-seq results, *CLDN4* was significantly down-regulated upon *ELF3* knockdown (p-value = 0.002254), and *CLDN7* also showed reduced expression, despite not meeting DE gene thresholds (p-value = 0.018557). No significant changes were observed in *CLDN3* and *EPCAM*. Unexpectedly, *VIM* expression was down-regulated (p-value = 0.003883), which contrasts with the reported role of *ELF3* as a suppressor of EMT and warrants further investigation. Replicate experiments confirmed these findings (**Supplementary Figure 9**).

Together, these results suggest that *ELF3* knockdown induces a partial Claudin-low phenotype in Basal-like cancer cells, as evidenced by the down-regulation of CLSD genes such as *CLDN4* and *CLDN7*.

## Discussion

In this study, we introduced a novel classifier UBS93 for the molecular subtyping of breast cancer. We validated its performance using both bulk and single-cell RNA sequencing data from human and mouse models. Notably, UBS93 outperformed classical methods, particularly in the identification of Claudin-low subtypes.

Using UBS93, we uncovered a previously unrecognized form of intratumoral heterogeneity in human Basal-like breast cancer, characterized by the co-existence of Basal-like and Claudin-low cancer cell populations within the same tumor. In our opinion, this finding carries three major implications. First, it advances our understanding of Basal-like breast cancer biology. Previously, such heterogeneity had only been reported in mouse models following genetic modulation of the RAS signaling pathway^5^. Whether this phenotype also occurs in humans remained unclear, given the substantial molecular differences between human and murine models. Our study provides direct evidence that this heterogeneity exists in human Basal-like breast tumors. Second, our findings offer compelling support for the hypothesis that Claudin-low cancer cells may originate from a Basal-like lineage, an idea previously proposed by Pommier et al. but lacking conclusive evidence. Third, our findings highlight the necessity of incorporating all intrinsic subtypes—including the Claudin-low subtype— into molecular subtyping frameworks, particularly in single-cell analyses. Notably, although Wu et al. introduced the SCSubtype classifier, its omission of the Claudin-low subtype led to the failure to detect this heterogeneity in human Basal-like scRNA-seq datasets.

Beyond the advancements in computational subtyping, our research has raised several intriguing scientific questions that warrant further investigation.

First, besides *ELF3* knockdown, what additional genetic perturbations are needed to fully convert Basal-like cancer cells into Claudin-low cancer cells? While high epithelial-to-mesenchymal transition (EMT) activity is a hallmark of Claudin-low cells, *ELF3* knockdown alone did not significantly up-regulate mesenchymal markers such as *VIM* or *FN1*. Interestingly, *ZEB1*—a key EMT regulator—was consistently up-regulated in the DE analyses comparing Claudin-low and Basal-like populations. This prompts the hypothesis that combined *ELF3* knockdown and *ZEB1* overexpression may be necessary to induce a complete Claudin-low phenotype.

Second, is *KRAS* mutation necessary for generation of Claudin-low cancer cells? Rädler *et al.* induced the tumor of “Basal-like/Claudin-low admixture” through activation of endogenous *KRAS* mutations (or exogenous *KRAS* expression). However, *KRAS* mutation is rare in human breast cancer. Using cBioPortal, we only identified six TCGA breast cancer patients carrying *KRAS* mutation; in addition, the cell line HDQP1 does not carry KRAS mutation either. These observations suggest that there might be *KRAS* independent mechanisms which may also give rise of Claudin-low cancer cells.

Third, what are the clinical implications of the “Basal-like/Claudin-low admixture”? Given that distant metastasis is the primary cause of cancer-related mortality and that Claudin-low tumors are associated with greater aggressiveness, it is plausible that the presence of such a phenotype may correlate with poor prognosis in Basal-like breast cancer. In future, we will integrate single-cell RNA-seq data with clinical outcome information to test this hypothesis.

Finally, does the presence of “Basal-like/Claudin-low admixture” alter the tumor microenvironment? Fougner *et al.* reported that Claudin-low tumors are characterized by high lymphocyte infiltration and elevated PD-L1 expression. We therefore hypothesize that Claudin-low cells within the tumors may shape distinct immune niches. To explore this, we will apply the UBS93 classifier to spatial transcriptomics datasets to identify regions enriched with Claudin-low cells and compare their surrounding microenvironments to those of Basal-like regions. This analysis may offer insights into immunotherapeutic vulnerabilities in this heterogeneous Basal-like breast cancer.

In conclusion, our study introduces a robust and versatile tool for delineating intratumoral heterogeneity in breast cancer. As high-quality single-cell transcriptomic data continue to emerge across diverse tumor types, we anticipate that this strategy will be broadly applicable to other cancers. By enabling precise dissection of tumor subpopulations, our approach holds promise for uncovering novel biological mechanisms that drive tumor progression and therapeutic resistance.

## Materials and Methods

### Development of UBS93 gene panel

We obtained a breast cancer scRNA-seq dataset consisting of 8 samples from Wu *et al.*’s research^16^. Given a gene g, its C-M-ratio value in sample S is calculated with the following formula:

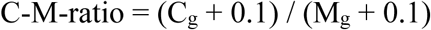

Here, C_g_ and M_g_ represent the average expression of G in cancer cells and non-cancer cells (of sample S), respectively.

Next, we divided all the samples into two groups: the TNBC group (2 samples) and the non-TNBC group (6 samples). We computed the group-wise C-M-ratio of gene g by taking the mean C-M-ratio within each group. Consequently, we obtained two group-wise C-M-ratio values for each gene g.

Finally, we ranked all genes according to their average group-wise C-M-ratio. We performed a quantile regression analysis between average group-wise C-M-ratio (Y) and rank (X) and noticed that there appears to be a change point of the slope. We picked out the top 400 genes to re-perform the analysis and found 300 appeared to be a reasonable estimate of the change point. Therefore, we finally kept the top 100 genes (**Supplementary Figure 10**). After excluding 7 genes that lack ENSEMBL gene IDs, we established the UBS93 gene panel.

### Determination of breast cancer subtypes

The CCLE categorized breast cancer into five subtypes: basal_A, basal_B, Luminal, HER2-amp, and luminal-HER2-amp. In the t-SNE plot (**Figure 1c**), basal_A and basal_B form their own distinct cluster while the other three subtypes form the third cluster. The basal_A cluster is significantly enriched with cell lines whose PAM50 subtype is Basal-like; therefore, it is reasonable to rename it as Basal-like; similarly, basal_B was renamed as Claudin-low. The luminal-HER2-amp subtype is not used in our classifier since the two cell lines of this subtype do not appear to form a distinct cluster.

### Differentiating Luminal and HER2-amp subtypes

We conducted differential gene expression analysis between CCLE HER2-amp and Luminal cell lines and identified 860 significantly up-regulated genes. We picked out the 20 genes which are present in the UBS93 panel and derived the HER2-amp signature.

In our classifier, ssGSEA analysis (UBS93 genes as background, and HER2-amp signature as a geneset) is utilized to differentiate Luminal and HER2-amp subtypes. We validated such strategy on a pooled bulk RNA-seq dataset (GSE212143 and GSE48213) and obtained an AUC (area under the curve) value of 0.964 (**Supplementary Figure 11**). In the final classifier, we determined the cutoff value to be 1.662 based on Youden’s index calculated in the validation dataset.

### Breast cancer subtyping with genefu

In genefu package, the gene sets used for Claudin-low and PAM50 subtyping are different. When analyzing a breast cancer sample, we first employ the function claudinLow to determine whether its subtype is Claudin-low; if not, we subsequently conduct PAM50 subtyping. In PAM50 subtyping, we removed the Normal-like subtype and merged LuminalA and LuminalB as Luminal.

### Gene set enrichment analysis

The EMT signature was downloaded from MSigDB^38^. To derive the Claudin-low-specific depletion (CLSD) gene set, z-score normalization was applied to gene expression values of CCLE breast cancer cell lines. For each gene, the median expression value across Claudin-low cell lines was computed. If this value is below -1, then the gene was considered as a CLSD gene. In total, this approach identified 960 CLSD genes. The clusterProfiler R package was used to compute enrichment score and p-value^39^.

### Processing of bulk RNA-seq data

We mapped raw bulk RNA-seq reads to transcriptome index with STAR^40^ and then utilized RSEM^41^ to quantify each gene’s TPM (Transcripts Per Million) value. The genome versions used are hg38 and mm10, respectively. The genome annotation files were downloaded from GENCODE^42^.

### The t-SNE analysis

Given two single cells C1 and C2, their distance is defined as 1-cor(C1, C2) where cor(C1, C2) is the Spearman rank correlation value across used genes. After the computation of the distance matrix, Rtsne function is used to generate the final embedding.

### Cell lines and cell culture conditions

HCC70 cells and HEK293T cells were obtained from Pricella Biotechnology Co., Ltd and American Type Culture Collection. HCC70 cells maintained in RPMI 1640 (catalog no.22400089, Gibco). HEK293T cells maintained in DMEM (catalog no. SH30285.01, Cytiva Life Sciences). All medium supplemented with 10% FBS (catalog no. 164210-50, Pricella Biotechnology).

### Lentivirus packaging and infection

HEK293T cells (2 × 10^6^) were seeded in a 100-mm cell culture dish and transfected with plasmids using PolyJet in vitro DNA transfection reagent PEI (C0537-20mg, Beyotime Biotechnology). The transfection experiments were operated according to the manufacturer’s instructions. For knockdown target proteins, we co-transfected the cells with the plasmids of pMDL, pVSVG, pREV and pLKO.1 (sh-*ELF3*, scrambled) to generate lentivirus. For lentivirus infection, 500 μl of virus-containing medium was added to the 5 × 10^4^ cells with 1% polybrene.

### Quantitative Real-time PCR

Total RNA was extracted from cells using an RNAsimple Total RNA Kit (TIANGEN, DP419). Complementary DNA was prepared using the High-Capacity cDNA Reverse Transcription Kit purchased from Vazyme Biotech (catalog no. R333–01). Real-time PCR analysis was performed using 2 × Universal SYBR Green Fast qPCR Mix (catalog no. RK21203, Abclonal Technology) with the following conditions: 3 min at 95 °C, and then 40 cycles at 95 °C for 5 s, 60 °C for 25 s, using a Real-Time System (Roche LightCycler® 480). Data were normalized according to expression of a control gene (*ACTB*) in each experiment. The data are presented as the mean ± s.d. from three independent experiments. The p-values were computed with unpaired t-test in GraphPad Prism 10 software.

### Single-cell RNA-sequencing of HCC70

The single-cell RNA-sequencing of HCC70 cell line was performed by a commercial company Novogene. Briefly, RNA from the barcoded cells was subsequently reverse-transcribed and sequencing libraries were constructed with reagents from a Chromium Single Cell 3’ v2 reagent kit (10X Genomics) according to the manufacturer’s instructions. Sequencing was performed with Illumina sequencing machines according to the manufacturer’s instructions. After filtering low-quality reads with fastp^43^, raw reads were demultiplexed and mapped to the human reference genome by cellranger 7.2.0^44^.

### Software tools and statistical methods

All the analysis was conducted in R. Seurat was used for processing scRNA-seq data. In DE analysis^45, 46, 47, 48^, genes with adjusted p-value < 0.05 and abs(log2FC) > 0.5) were considered as differentially expressed genes. The ggplot2 package was used for data visualization, pheatmap and ComplexHeatmap packages were used to generate heat maps^49,50,51^, GSVA was used for ssGSEA analysis^52, 53^. If not specified, the two-sided Wilcoxon rank-sum test was used to compute the p-value in hypothesis testing.

## Data Availability

The authors declare that all data supporting the findings of this study are available within the article and its supplementary information files or from the corresponding author upon reasonable request.

## Code Availability

The code is available at github (https://github.com/bioklab/UBS93).

## Acknowledgments

We thank Dr. Xiaohui Lin for helping us revise the manuscript. The research is supported by National Natural Science Foundation of China (Fund 32370715), Science Fund for Distinguished Oversea Young Scholars of Shandong Province (2023HWYQ-015), Taishan Young Scholar Program of Shandong Province (tsqn202312020), and Cheeloo Young Scholar Program of Shandong University. The content is solely the responsibility of the authors and does not necessarily represent the official views of sponsors.

## Authors’ contribution

J.L. and K.L. conceived the study. J.L. performed the majority of computational analysis, Y.G. and W.X.Y. performed all the experimental assays, and K.L. supervised the study. All authors contributed to writing, reviewing, and editing the manuscript and approved the manuscript.

