## Supplementary material for "Single-cell molecular subtyping reveals novel intratumor heterogeneity in human Basal-like breast cancer": Supp Figure

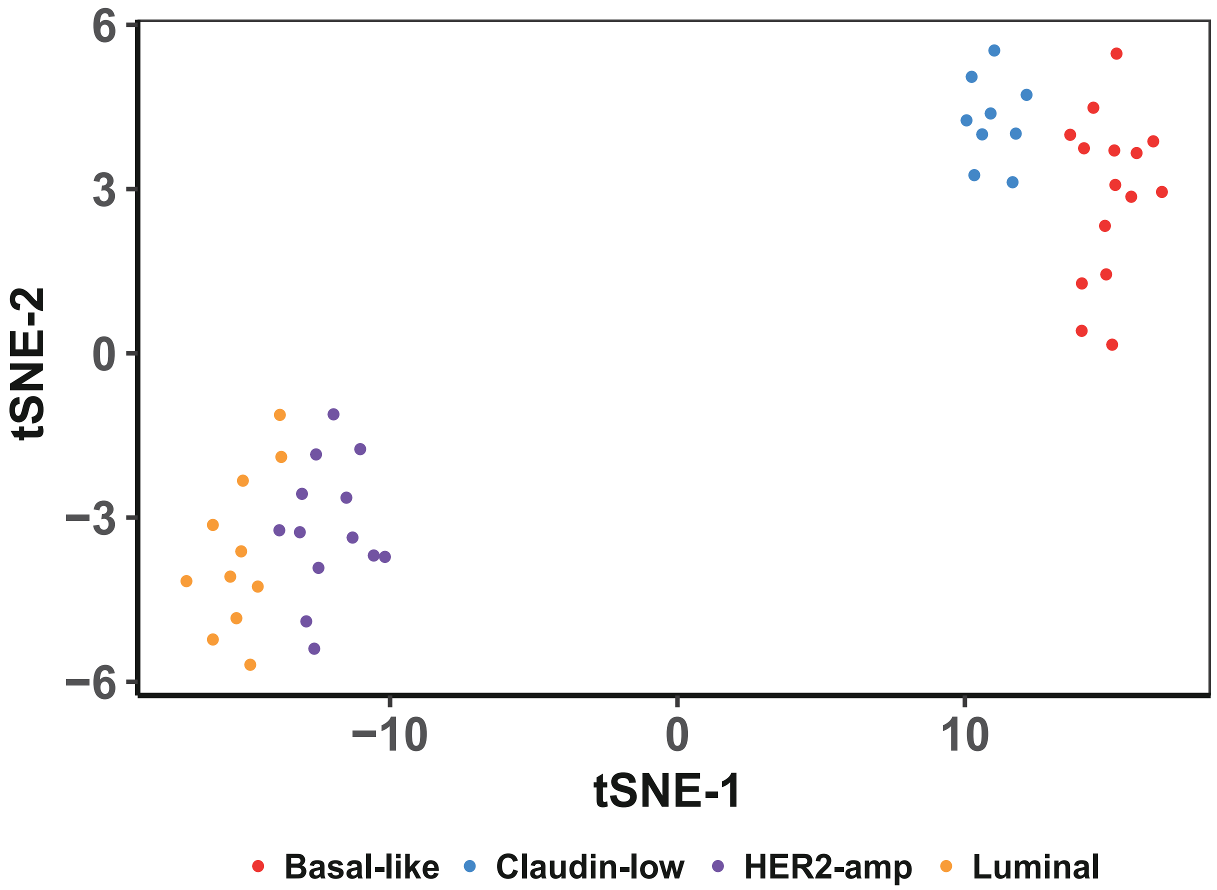


**Supplementary Figure 1:** The t-SNE visualization of CCLE breast cancer cell lines based on the UBS93 gene panel.

a

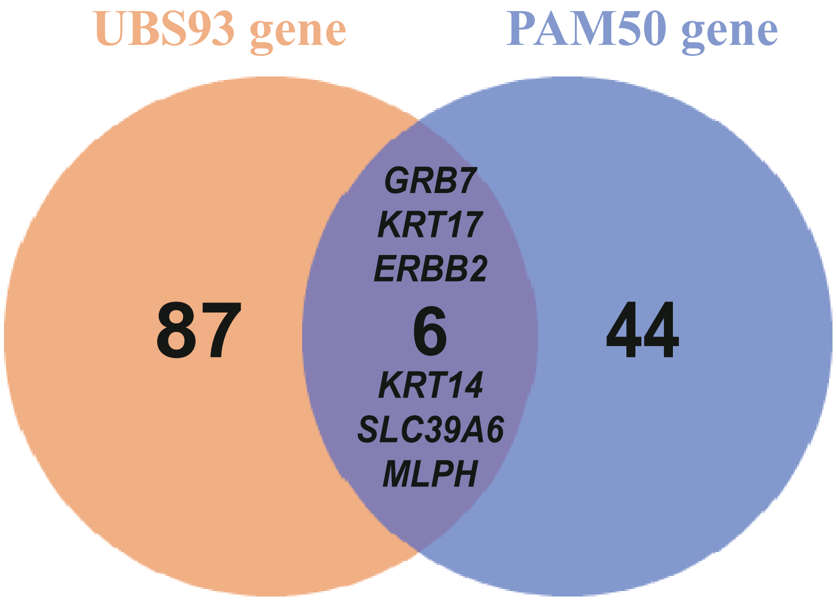


b

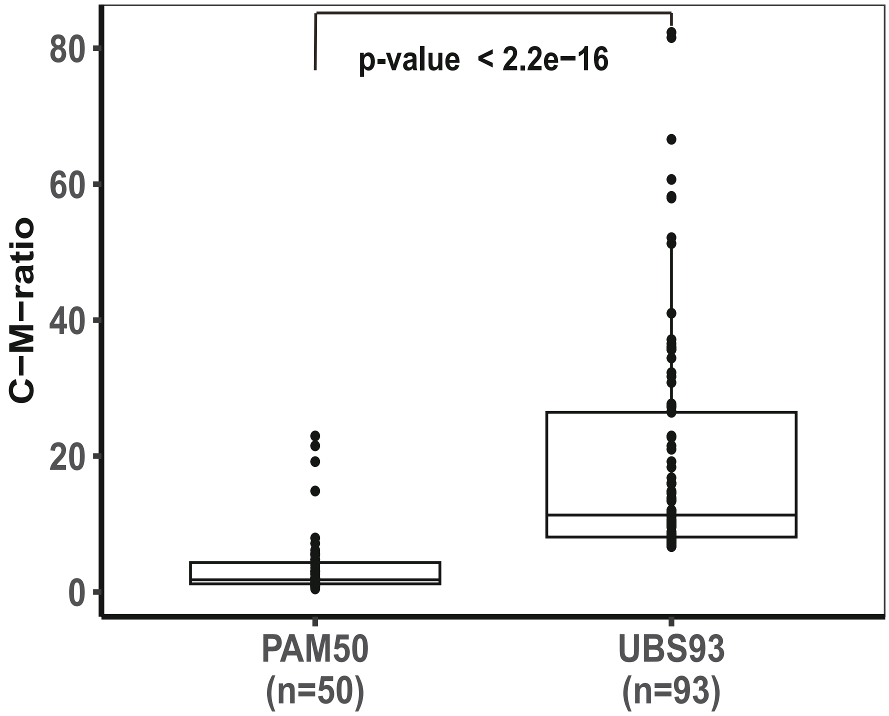


**Supplementary Figure 2:**
(a) Venn diagram comparing UBS93 and PAM50 genes.
(b) C-M ratios of PAM50 and UBS93 genes. Each dot represents a gene, the central line indicates the median, and the bounds represent the 25th and 75th percentiles (interquartile range). Whiskers extend to 1.5 times the interquartile range. P-values were calculated using the two-sided Wilcoxon rank-sum test.

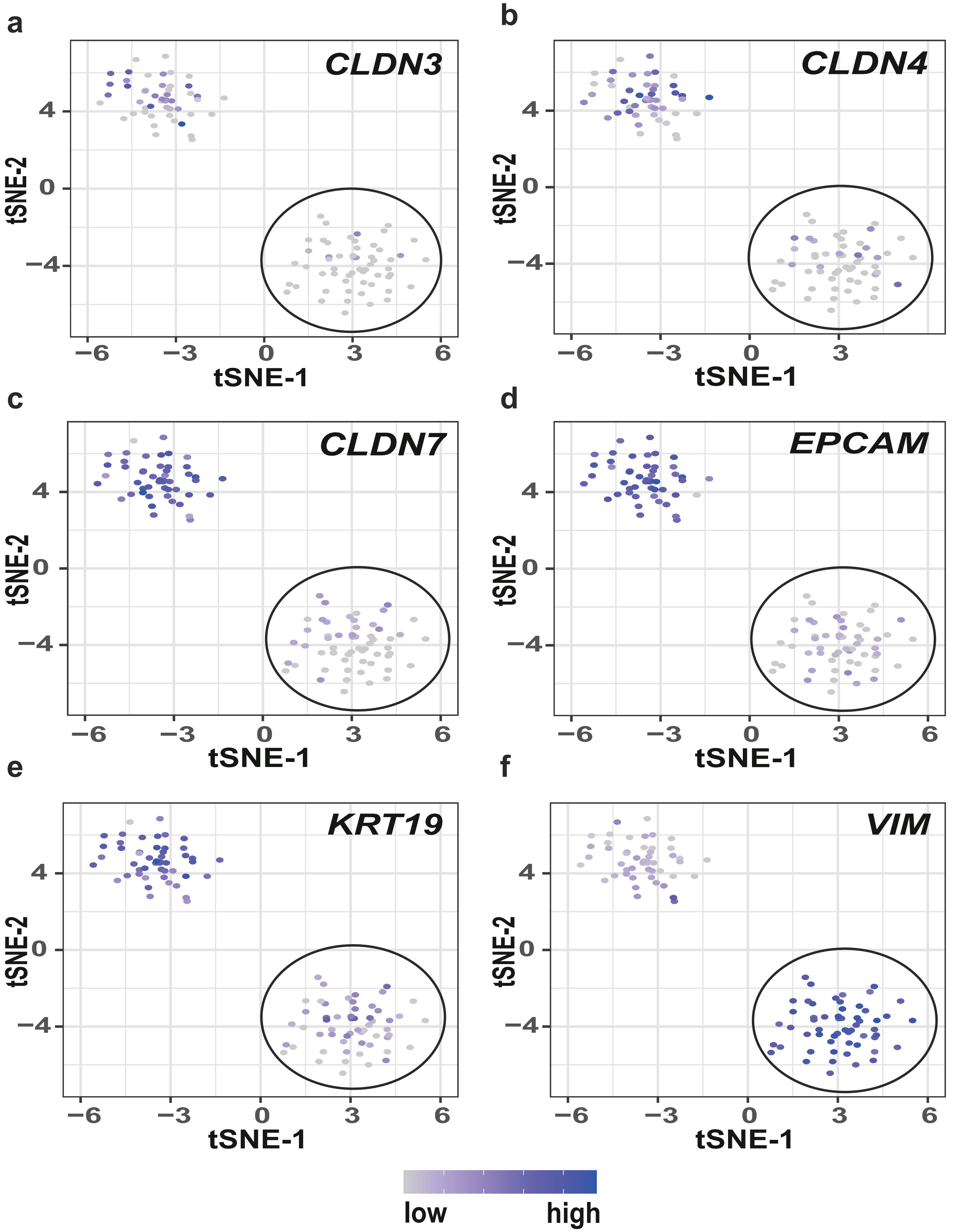


**Supplementary Figure 3:** Expression of *CLDN3* (a), *CLDN4* (b), *CLDN7* (c), *EPCAM* (d), *KRT19* (e), and *VIM* (f) in HDQP1 single cells (from GSE173634 dataset). The circled region highlights a cluster where the majority of cells belong to the Claudin-low subtype.

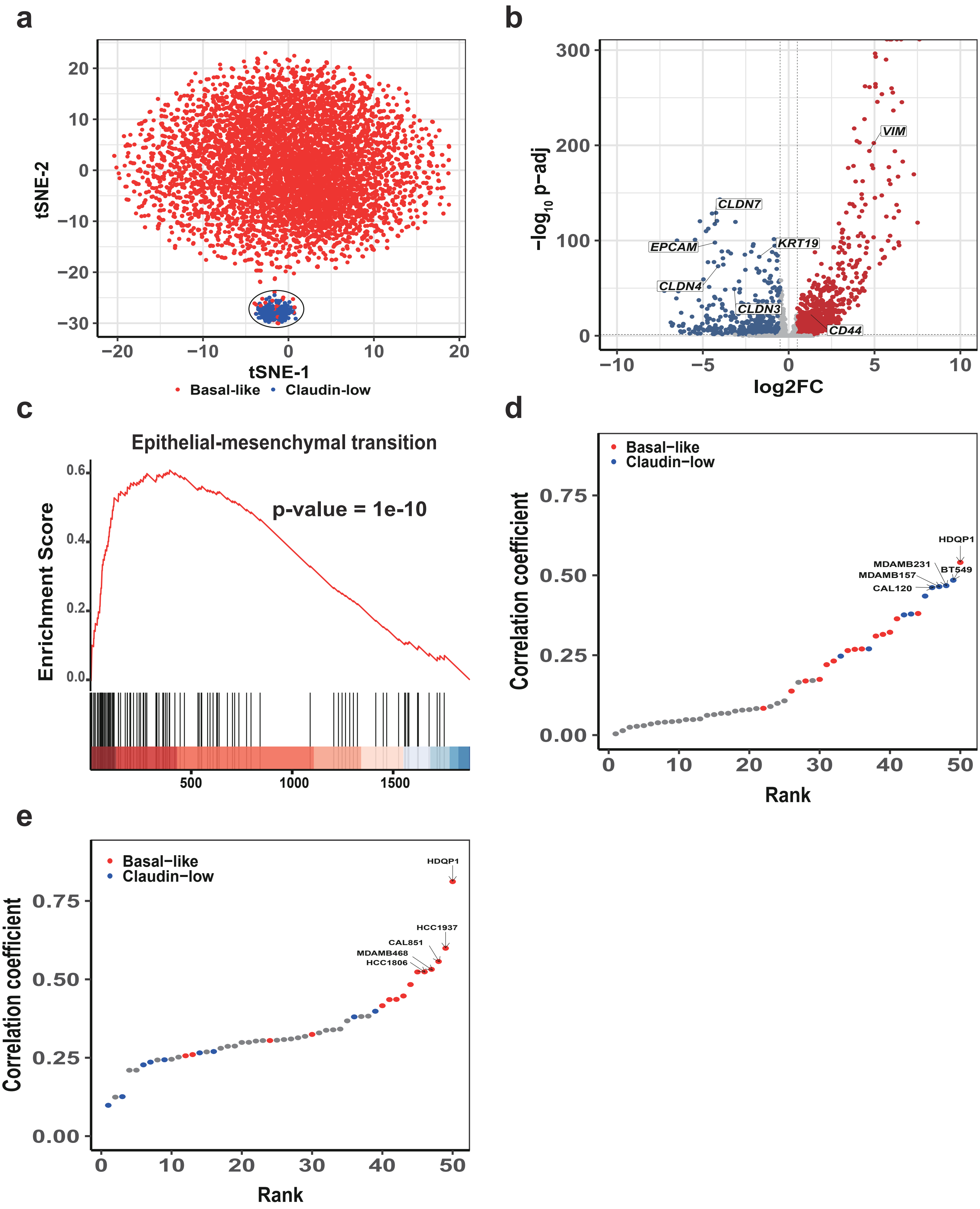
a
tSNE-2
tSNE-1
Basal-like
Claudin-low
b
−log10 p−adj
log2FC
CLDN7
EPCAM
CLDN4
CLDN3
KRT19
VIM
CD44
c
Epithelial-mesenchymal transition
Enrichment Score
p-value = 1e-10
d
Basal−like
Claudin−low
Correlation coefficient
Rank
HDQP1
MDAMB231
MDAMB157
CAL120
BT549
e
Basal−like
Claudin−low
Correlation coefficient
Rank
HDQP1
HCC1937
CAL851
MDAMB468
HCC1806

**Supplementary Figure 4:**
(a) The t-SNE visualization of HDQP1 scRNA-seq data (GSE202771), with cells colored by UBS93 subtypes. The circled region highlights a cluster where the majority of cells belong to the Claudin-low subtype.
(b) Differential gene expression analysis comparing Claudin-low and Basal-like cells of the HDQP1 cell line. Red and blue dots represent genes significantly up-regulated and down-regulated in Claudin-low cells, respectively.
(c) Gene set enrichment analysis (GSEA) of epithelial-mesenchymal transition (EMT) signature in the differentially expressed genes. The barcode beneath the heatmap represents the ranked position of EMT genes, from red (high log2FC) to blue (low log2FC).
(d-e) Ranking of CCLE breast cancer cell lines based on transcriptomic correlation with Claudin-low (or Basal-like) pseudo-bulk sample. The Basal-like and Claudin-low cell lines are colored red and blue, respectively.

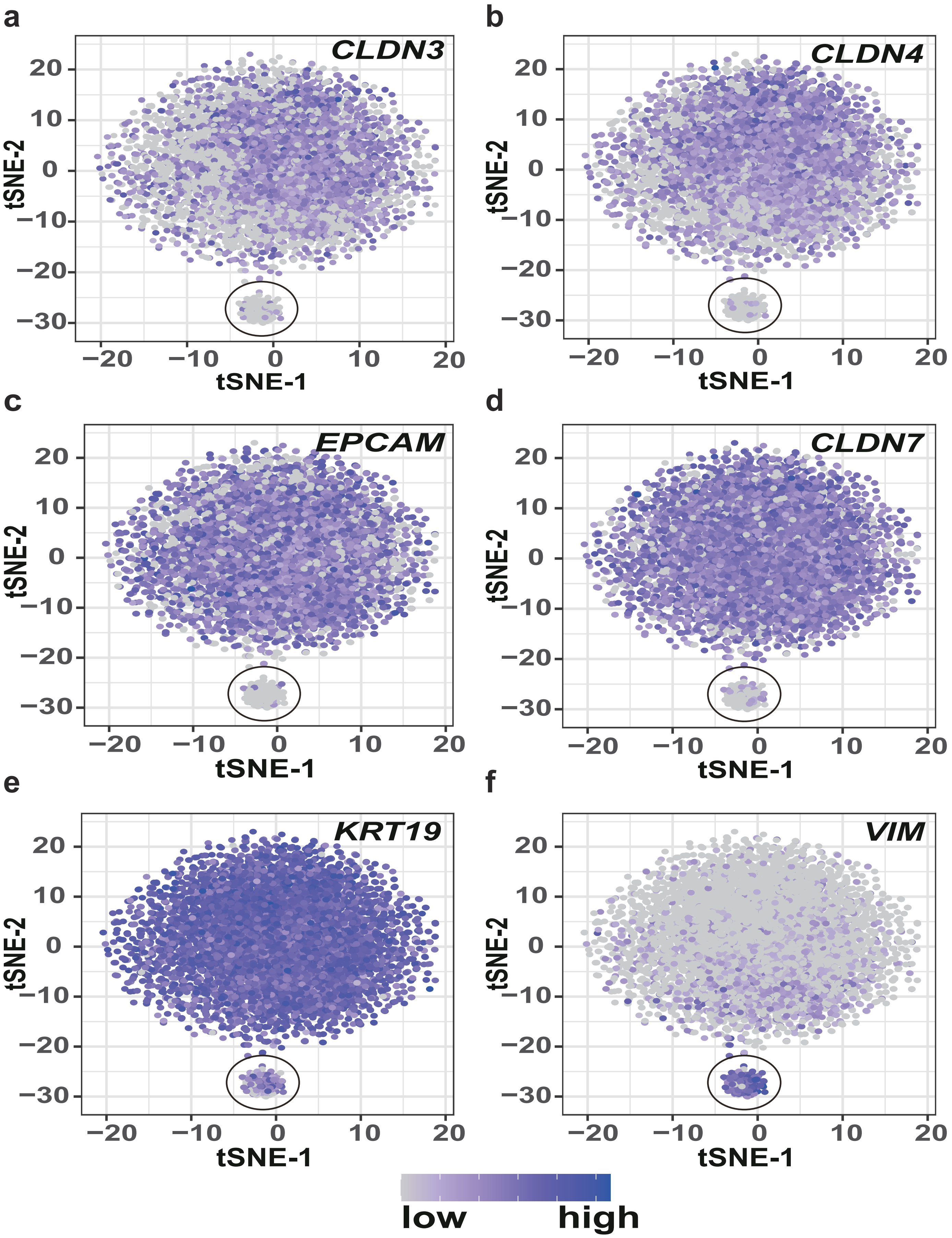


**Supplementary Figure 5:** Expression of *CLDN3* (a), *CLDN4* (b), *CLDN7* (c), *EPCAM* (d), *KRT19* (e), and *VIM* (f) in HDQP1 single cells (from GSE202771 dataset). The circled region highlights a cluster where the majority of cells belong to the Claudin-low subtype.

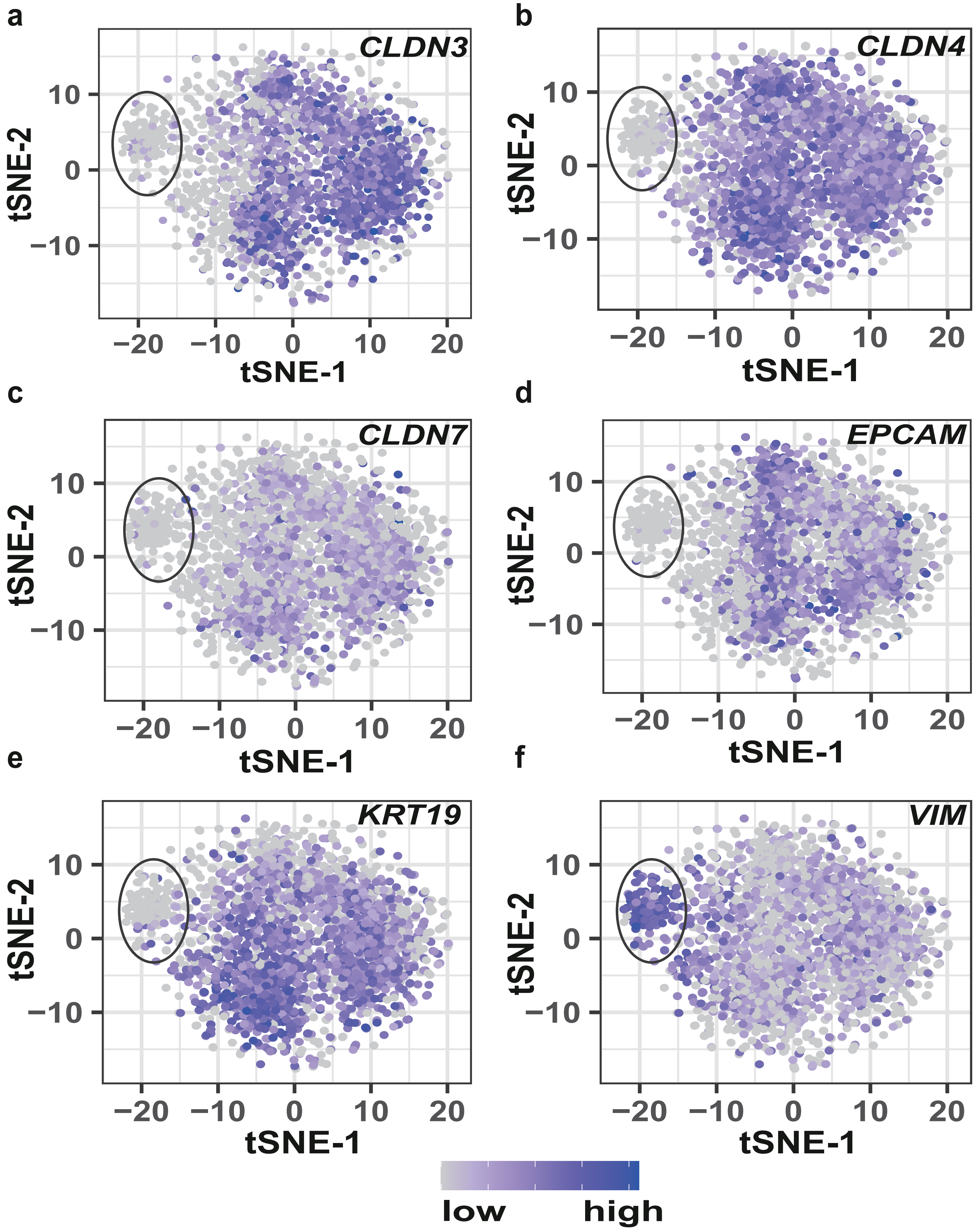


**Supplementary Figure 6:** Expression of *CLDN3* (a), *CLDN4* (b), *CLDN7* (c), *EPCAM* (d), *KRT19* (e), and *VIM* (f) in cancer cells from TNBC patient 0554. The circled region highlights a cluster where the majority of cells belong to the Claudin-low subtype.

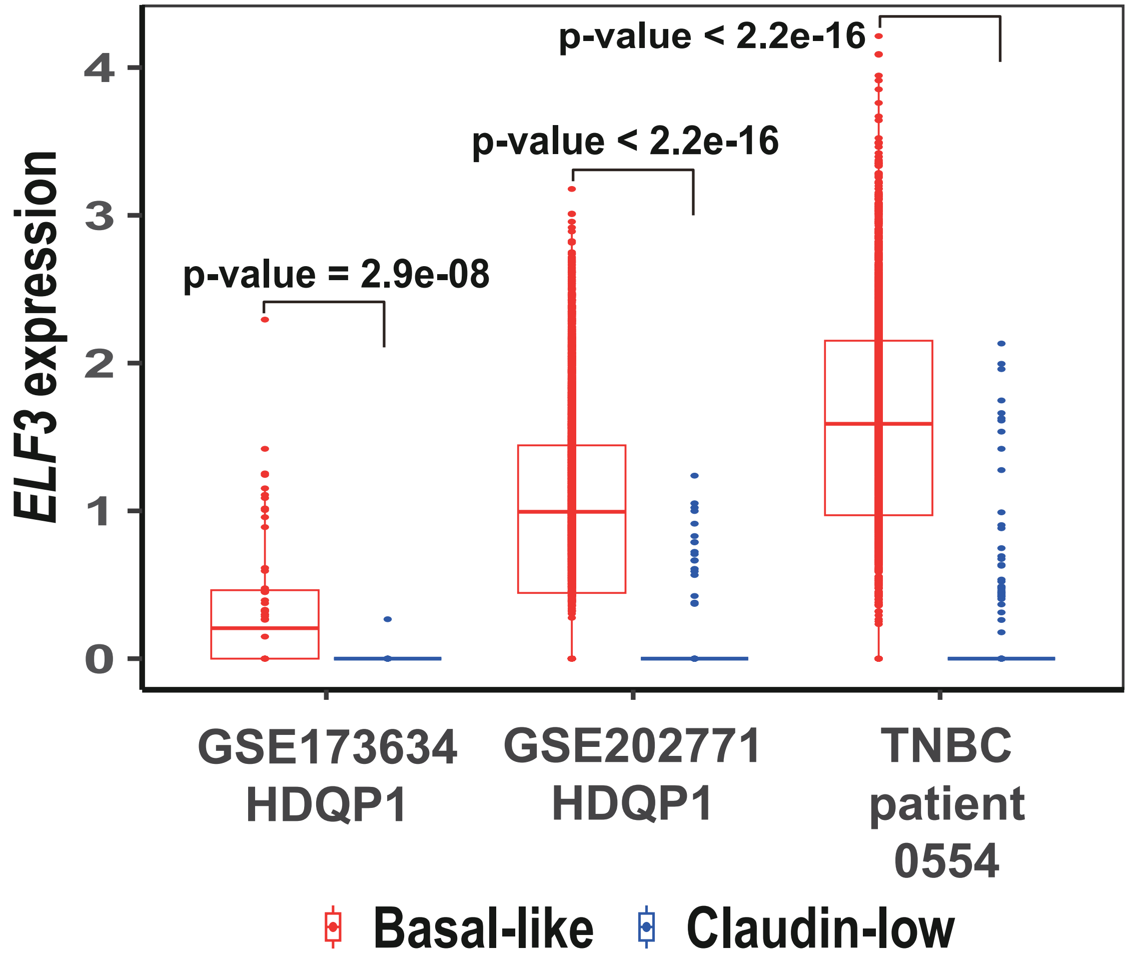


**Supplementary Figure 7:** *ELF3* expression in Basal-like and Claudin-low cancer cells in HDQP1 cell line (from GSE173634 and GSE202771 datasets) and TNBC patient 0554.

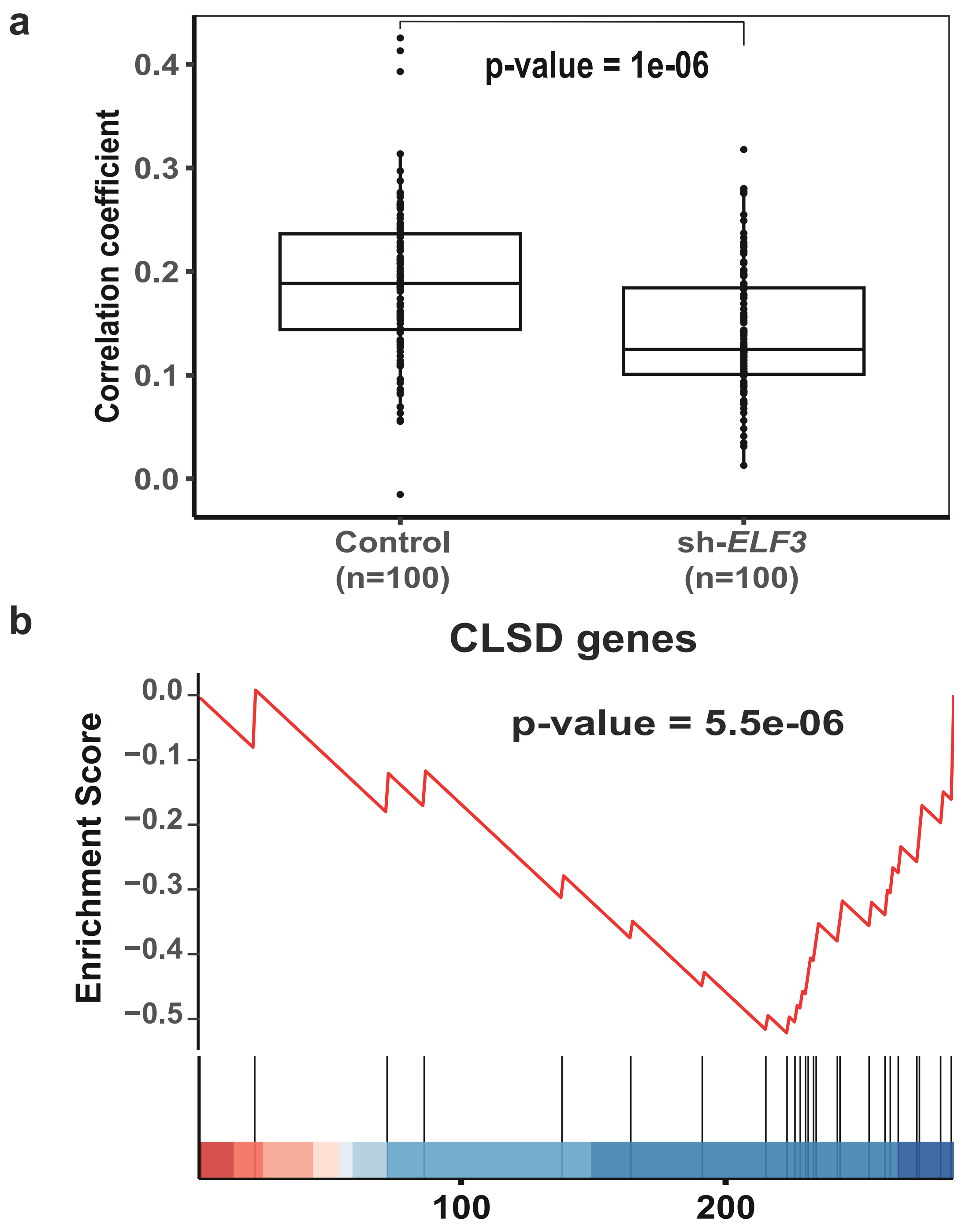


**Supplementary Figure 8:**
(a) Correlation coefficients between the Basal-like centroid and the two groups of single cells (control and sh-*ELF3*). Each dot represents a single cell, the central line indicates the median, and the bounds represent the 25th and 75th percentiles (interquartile range). Whiskers extend to 1.5 times the interquartile range. P-values were calculated using the two-sided Wilcoxon rank-sum test.
(b) Gene set enrichment analysis (GSEA) of CLSD gene signature in differentially expressed genes (sh-*ELF3* vs. control). The barcode beneath the heatmap represents the ranked position of genes from red (high log2FC) to blue (low log2FC).

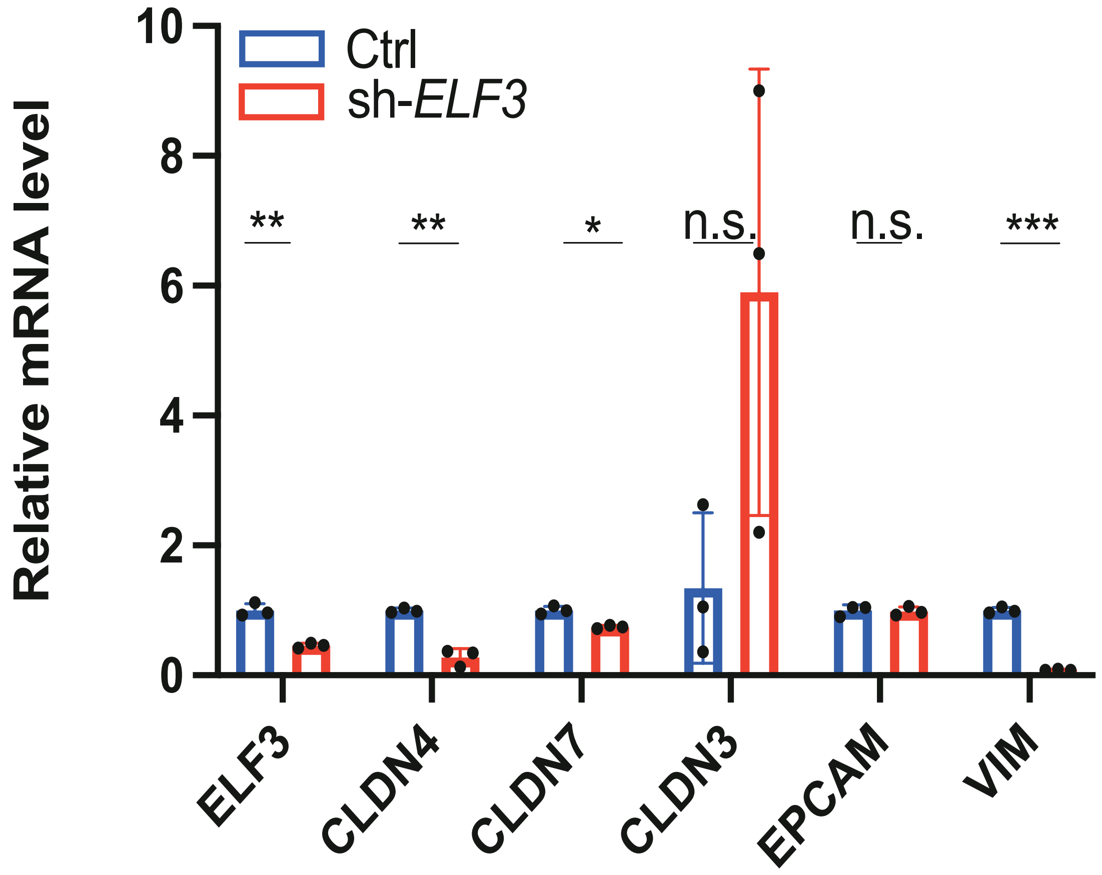


**Supplementary Figure 9:** Relative mRNA levels of *ELF3*, *CLDN4*, *CLDN7*, *CLDN3*, *EPCAM*, and *VIM* in the control group (blue) and the sh-*ELF3* group (red). Each dot represents a parallel sample. P-values were calculated using the two-sided t-test. n.s. ($p > 0.05$), *($p < 0.05$), ** ($p < 0.01$), *** ($p < 0.001$).

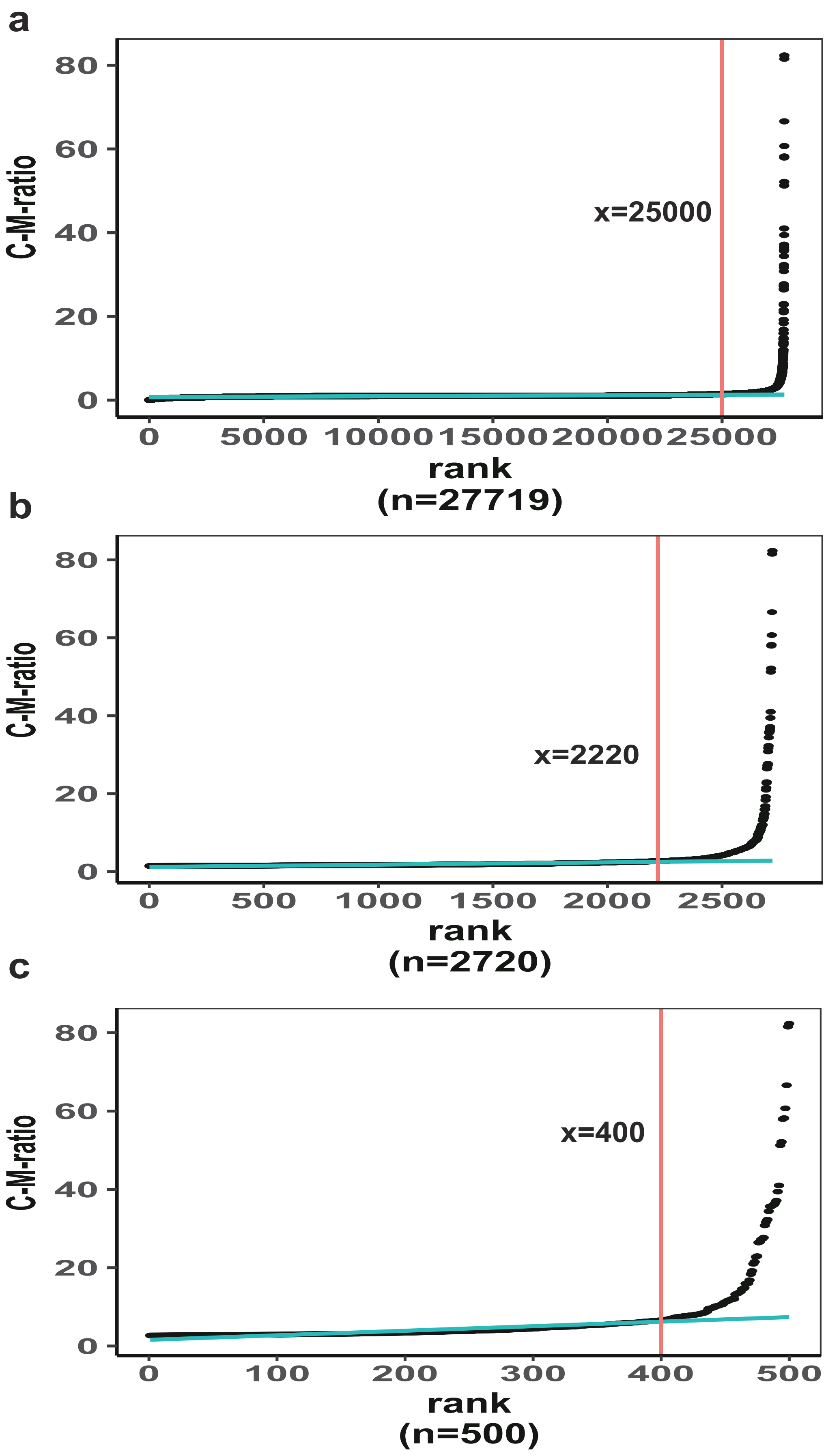


**Supplementary Figure 10:** Using quantile regressions to determine number of genes with high C-M-ratio values. In each panel, the vertical red line indicates a manually determined threshold of rank, and genes (in panel a anb b) whose rank values are higher than the threshold are further used in the next panel. The cyan line represents the fitted regression line.

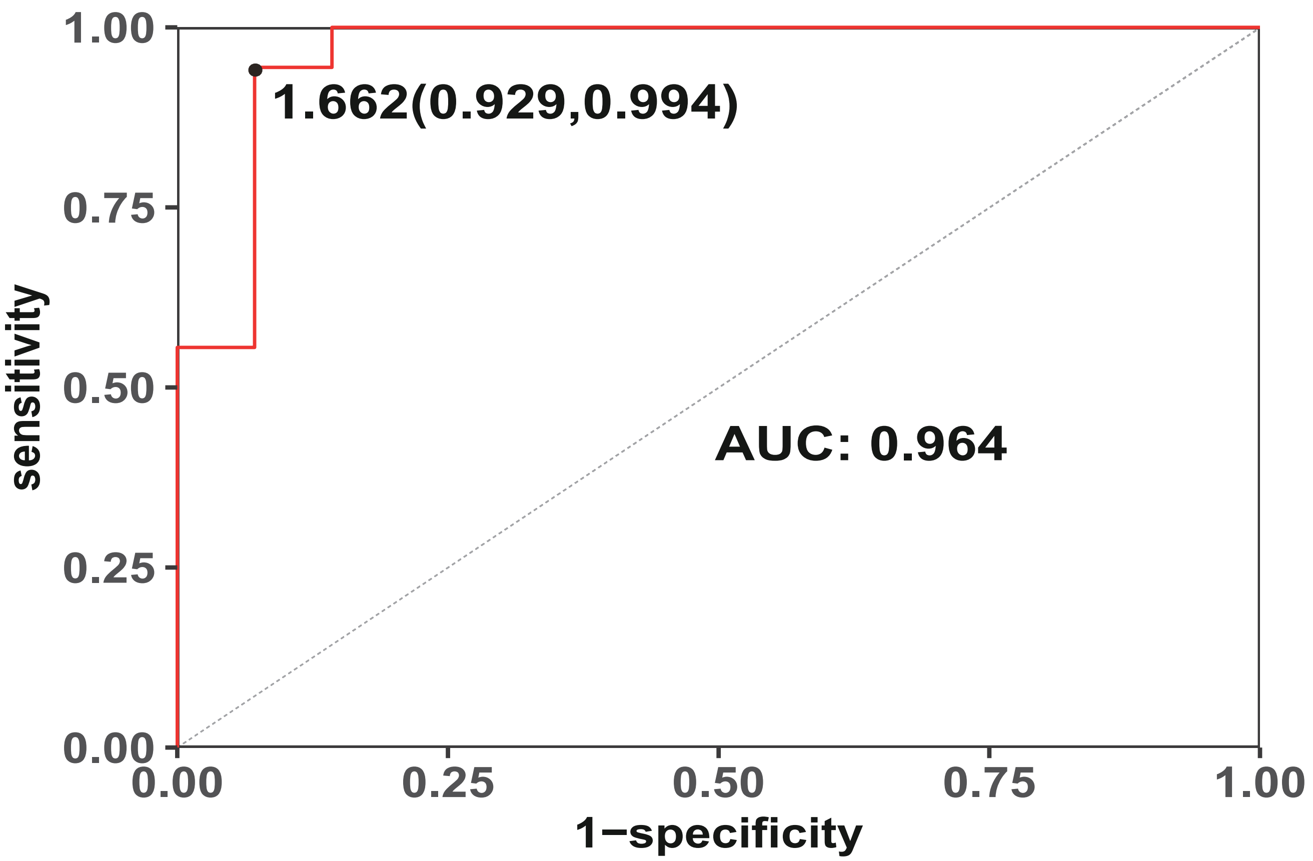


**Supplementary Figure 11:** Differentiating Luminal and HER2-amp subtypes using ssGSEA score. The ROC curve was plotted using a combined dataset (GSE212143 and GSE48213) and the Youden index was highlighted.
